# Motor and Sensory Cortical Processing of Neural Oscillatory Activities revealed by Human Swallowing using Intracranial Electrodes

**DOI:** 10.1101/2020.07.21.213868

**Authors:** Hiroaki Hashimoto, Kazutaka Takahashi, Seiji Kameda, Fumiaki Yoshida, Hitoshi Maezawa, Satoru Oshino, Naoki Tani, Hui Ming Khoo, Takufumi Yanagisawa, Toshiki Yoshimine, Haruhiko Kishima, Masayuki Hirata

**Affiliations:** Department of Neurological Diagnosis and Restoration, Graduate School of Medicine, Osaka University, Yamadaoka 2-2, Suita, Osaka 565-0871, Japan; Department of Neurosurgery, Otemae Hospital, Chuo-ku Otemae 1-5-34, Osaka, Osaka 540-0008, Japan; Endowed Research Department of Clinical Neuroengineering, Global Center for Medical Engineering and Informatics, Osaka University, Yamadaoka 2-2, Suita, Osaka 565-0871, Japan; Department of Organismal Biology and Anatomy, University of Chicago, 1027 E 57th St, Chicago, IL 60637, USA; Department of Anatomy and Physiology, Saga Medical School Faculty of Medicine, Saga University, Nabeshima 5-1-1, Saga, Saga 849-8501, Japan; Department of Neurosurgery, Graduate School of Medicine, Osaka University, Yamadaoka 2-2, Suita, Osaka 565-0871, Japan

## Abstract

Swallowing, a unique movement, is attributed to the indispensable orchestration of motor-output and sensory-input. We hypothesized that swallowing can illustrate differences between motor and sensory neural processing. Eight epileptic participants fitted with intracranial electrodes over the orofacial cortex were asked to swallow a water bolus. Mouth-opening and swallowing were treated as motor tasks, while water-injection as sensory tasks. Phase-amplitude coupling between lower frequency and high γ (HG) band (75–150 Hz) was investigated. An α (10-16 Hz) -HG coupling appeared before motor-related HG power increase (burst), and a θ (5-9 Hz) -HG coupling appeared during sensory-related HG burst. The motor- and sensory-related HG amplitude were modulated at the trough of α oscillations and peak of θ oscillations, respectively. These contrasting results acquired from the orofacial cortex can help to fully elucidate the sensory-motor function in the brain.

## Introduction

Motor- and sensory-related tasks induce neural oscillatory changes in the cortex. Scalp electroencephalogram (EEG) studies have demonstrated that finger movement induced power suppression in the α band, representing mu activity (Pfurtscheller and Aranibar, 1979). During movement or stimulation task activating sensorimotor cortex, power suppression of beta band was also observed (Pfurtscheller, 1981). Oscillatory changes have been reported using sensory stimulation in a magnetoencephalography (MEG) study (Hirata et al., 2002, Maezawa et al., 2008). Suppressed β bands are re-activated following the completion of motor task, which is referred to as the “rebound β” (Salmelin and Sams, 2002). These frequency bands, i.e., the α (8-14 Hz) (Haegens et al., 2011) and β (15-25 Hz) (Crone et al., 1998b), are lower frequency band (Miller et al., 2007), whereas, frequencies over 30 Hz are γ bands, and those > 75 Hz are referred to as high γ (HG) band (Crone et al., 1998a).

HG activities are involved in not only sensory (Hirata et al., 2002) and motor tasks (Crone et al., 1998a, Dalal et al., 2008), but also attention (Taylor et al., 2005), language (Hashimoto et al., 2017), and working memory (Pipa et al., 2009). HG activity is a key oscillation that reflects the neural processing and shows better functional localization than lower frequency activity (Crone et al., 1998a, Canolty et al., 2006, Miller et al., 2007). These high frequency activities (HG activities) are physiological activities; pathological high frequency activities have been previously reported in epilepsy (Matsumoto et al., 2013) and are referred to as high frequency oscillations (Jacobs et al., 2009) or high frequency activities (HFA) (Ayoubian et al., 2013).

The amplitude of HG activities is modulated by a phase of low-frequency oscillations (Canolty et al., 2006). The relationship between the HG amplitude and the phase of low-frequency oscillations has been investigated using phase-amplitude coupling (PAC) methods (Cohen, 2008). PAC plays different functional roles in cortical processing, such as motor execution (Yanagisawa et al., 2012b) and somatosensory processing (Lakatos et al., 2008). These cross-frequency coupling are physiological neural processing, while pathological cross-frequency coupling have also been reported to be associated with epileptic seizures (Hashimoto et al., 2020b, Hashimoto et al., 2020a), and Parkinson disease (De Hemptinne et al., 2015b).

The above neural oscillations were investigated for clinical applications. The lower frequency neural oscillations in sensorimotor cortex measured by scalp EEG were used for decoding in rehabilitation using brain-machine interface (BMI) techniques (Ramos-Murguialday et al., 2013, Shindo et al., 2011). Whereas HG activities could be used to decode more accurate, rather than lower, frequency bands (Yanagisawa et al., 2011, Yanagisawa et al., 2012a, Hashimoto et al.). Cross-frequency coupling could be used to detected or predicted epileptic seizures (Amiri et al., 2019, Edakawa et al., 2016).

To investigate motor- and sensory-related neural oscillations, various tasks have been used including motor-related tasks include button pressing (Pfurtscheller and Aranibar, 1979), hand and elbow movements (Yanagisawa et al., 2012a), and tongue movements (Miller et al., 2007) as well as sensory-related tasks include a median nerve stimulation (Hirata et al., 2002), a tongue stimulation (Maezawa et al., 2008), and a buccal mucosa stimulation (Miyaji et al., 2014). Even swallowing is used as a task to investigate neural oscillatory changes (Cuellar et al., 2016, Dziewas et al., 2003, Furlong et al., 2004, Satow et al., 2004). We have previously reported that swallowing-related HG activities were observed in the cortex along the Sylvian fissure (Hashimoto et al., 2020c).

Swallowing is a unique movement since cooperation between motor output and sensory input are involved. Sensory input is essential to the initiation and modulation of normal swallowing (Gow et al., 2004, Lowell et al., 2008); anesthesia of the oral and pharynx area can interfere with swallowing (MÅrnsson and Sandberg, 1974). Moreover, various muscles of the oral, pharynx, and esophagus areas are involved in swallowing execution (Ertekin and Aydogdu, 2003). Swallowing water task has two aspects: A motor-output aspect that includes mouth open and closely followed by swallowing, while the other is a sensory-input aspect that includes stimulation of a buccal mucosa by water injection into the oral cavity. Therefore, we hypothesized that swallowing water task can show differences between motor and sensory neural processing.

This study has two aims: One, since the role of PAC in swallowing remains unknown, to determine whether PAC is observed during the swallowing task in the orofacial cortex, and two, to identify the differences between motor- and sensory-related neural processing using a combination of swallowing water task and PAC methods.

Neurophysiological recording techniques, such as the intracranial EEG (iEEG), or electrocorticogram (ECoG), can identify neural oscillatory changes up to HG bands. In this study, we recorded iEEG signals in the orofacial cortex of patients with epilepsy while swallowing a bolus of water. We continue to research for contributions to recovery from dysphagia using swallowing-related neural activities in combination with our BMI technology. As a part of this research, we have previously reported the development of a swallow-tracking system that enabled us to monitor swallowing noninvasively using Kinect v2 (Microsoft, Redmond, Washington, USA) (Hashimoto et al., 2018). We can use this swallow-tracking system simultaneously with iEEG measurement to detect the time when participants open their mouth, when water bolus were injected into the mouth, and when participants swallowed. Therefore, we could insert three different triggers into iEEG signals; mouth-triggers (Mouth-T), water-triggers (Water-T), and swallow-triggers (Swallow-T) which corresponded to the three different times, respectively. We have also analyzed the iEEG signals that are time-locked to the triggers. Mouth-T and Swallow-T represented a motor-task, while Water-T represented a sensory-task. The synchronization index (SI) was used for PAC analysis (Cohen, 2008).

## Results

### Distinct high γ activities were evoked by mouth opening, water injection, and swallowing

For a representative participant (Participant 1; P1), the spatial HG normalized power distributions in the orofacial cortex are shown in Figure 1A. Within three different triggers including Mouth-T, Water-T, and Swallow-T, significant HG power increasing contacts (corrected *p* (*c.p*) < 0.05, single-sided permutation tests with Bonferroni correction) are indicated as filled white circles, and we could observe a different spatial distribution.

**Fig.1.**
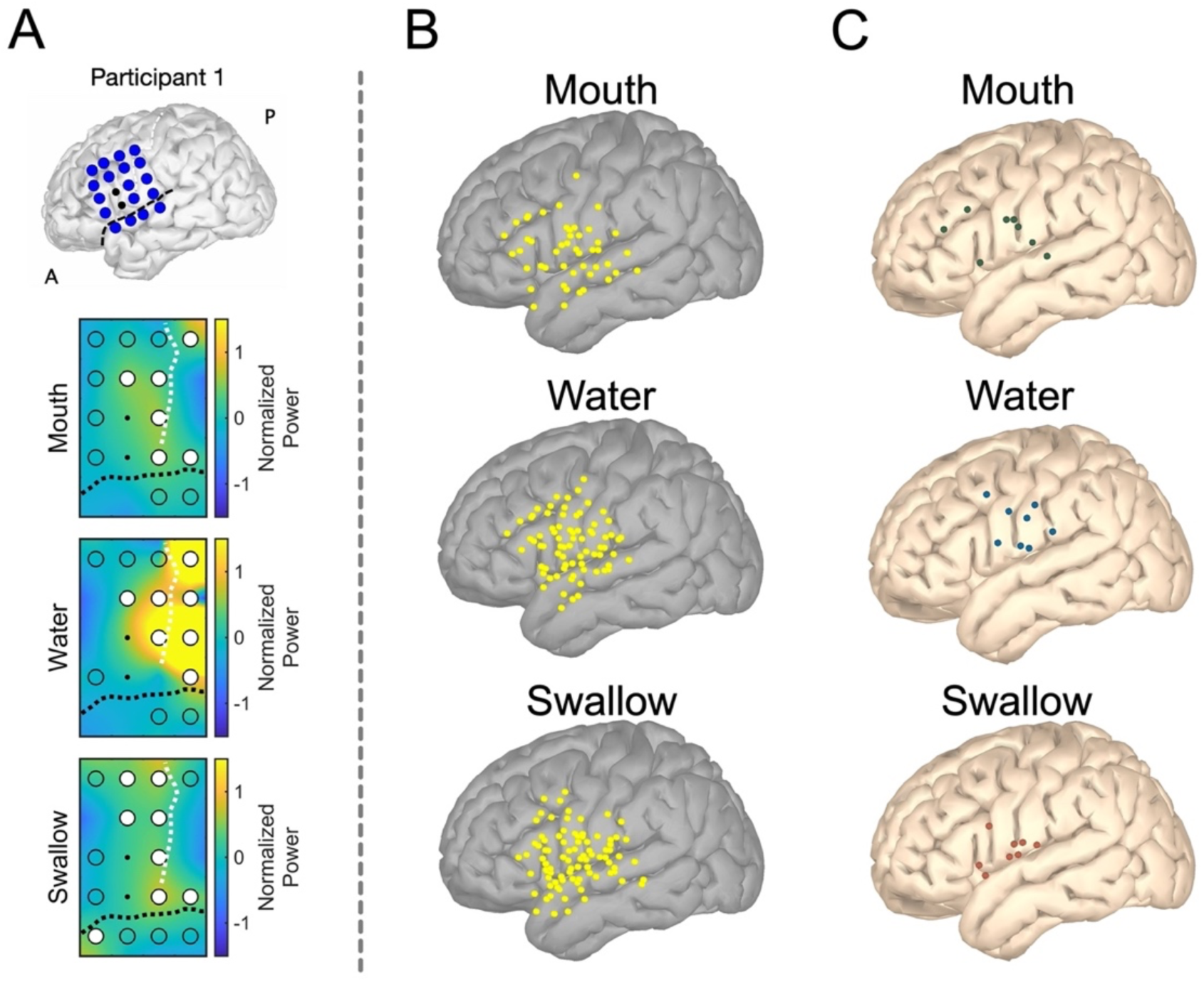
Distributions of high γ (HG) activities in the orofacial cortex. (A) The reconstructed magnetic resonance images of Participant 1 are demonstrated at the top. The central sulcus and the Sylvian fissure are indicated by white and black dashed lines, respectively. A; anterior, P; posterior. Contacts of implanted intracranial electrodes are indicated by blue circles. The two black contacts were excluded due to severe noise contamination. The contour maps indicate distributions of HG normalize power within mouth, water, and swallow triggers. The pattern of HG distributions varies among three triggers. Significant power increasing is indicated as white filled circles (corrected *p* < 0.05, single-sided permutation test with Bonferroni correction). (B) For each trigger group, contacts indicating significant power increases in HG band of all participants are plotted over the left hemisphere of the Montreal Neurological Institute (MNI) brain. (C) The contacts indicating the maximum values of significant HG power increase from each participant are plotted on the left hemisphere of MNI normalized brain. Within each trigger, the contacts group are defined as a mouth contacts group (Mouth-C), a water contacts group (Water-C), and a swallow contacts group (Swallow-C), colored by green, blue, and red in each.

Total eight participants were enrolled (Table 1). All contacts showing HG power increase in all participants were plotted on the left hemisphere of the Montreal Neurological Institute (MNI) normalized brain (Fig. 1B). For better distributions clarity, we selected the contacts indicating the maximum values of significant HG power increase within Mouth-T, Water-T, and Swallow-T from each participant, and the total eight contacts within each trigger were plotted on the left hemisphere of MNI normalized brain (Fig. 1C). With Mouth-T, HG increasing was observed primarily in the frontal lobe, and with Water-T, HG increasing was observed in the lateral pericentral gyri. With Swallow-T, HG increasing appeared in the region along the Sylvian fissure. These three contacts-groups were defined as a mouth contacts group (Mouth-C), a water contacts group (Water-C), and a swallow contacts group (Swallow-C).

**Table 1.** Clinical profiles. M, Male; F, Female; R, Right; L, Left

| Participant | Sex | Age | Side | Number of orofacial contacts (Total contacts) | number of swallowing instances | number of mouth openings | Number of water injections |
| --- | --- | --- | --- | --- | --- | --- | --- |
| P1 | F | 30's | L | 18 (68) | 31 | 31 | 31 |
| P2 | F | 30's | L | 20 (84) | 41 | 41 | 41 |
| P3 | F | 10's | R | 18 (55) | 27 | 27 | 27 |
| P4 | M | 20's | L | 15 (69) | 34 | 34 | 34 |
| P5 | M | 50's | L | 20 (72) | 38 | 38 | 38 |
| P6 | M | 20's | R | 20 (96) | 27 | 27 | 27 |
| P7 | M | 20's | L | 20 (94) | 33 | 33 | 33 |
| P8 | F | 10's | L | 20 (68) | 37 | 37 | 37 |

Averaged normalized power of HG around Mouth-T, Water-T, and Swallow-T were calculated from Mouth-C, Water-C, and Swallow-C (Fig. 2A). Around 0 s of Mouth-T, Water-T, and Swallow-T for each, notable HG power burst were observed in Mouth-C, Water-C, and Swallow-C respectively.

**Fig.2.**
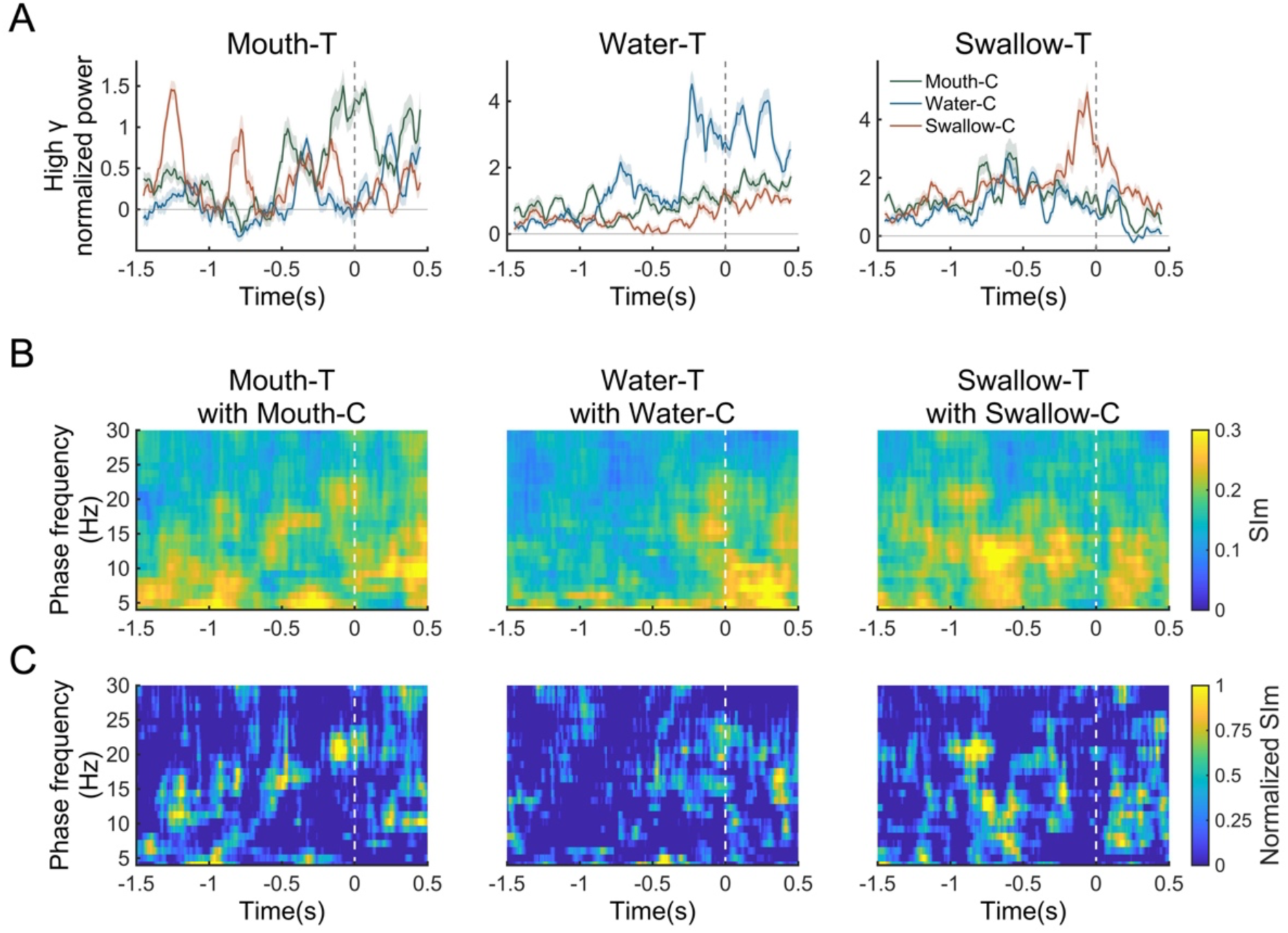
Temporal profile of high γ (HG) activities and phase-amplitude coupling (PAC). (A) Averaged normalized power of HG calculated from Mouth-C, Water-C, and Swallow-C are plotted from −1.5 s to 0.5 s around Mouth-T, Water-T, and Swallow-T. At 0 s of Mouth T, only Mouth-C showed a HG burst not Water-C and Swallow-C. At 0 s of Water T, Water-C showed notably HG bursts. With Swallow T, Swallow-C showed HG power increasing not Mouth-C and Water-C. The error bars indicate 95% confidence intervals. (B) For PAC analyses, the magnitudes of synchronization index (SIm) were calculated between lower frequency (5-30 Hz, 1 Hz bin) phase (vertical axes) and HG (75-150 Hz) amplitude. We used Mouth-C in Mouth-T, Water-C in Water-T, and Swallow-C in Swallow-T. In motor-related PAC including Mouth-T with Mouth-C and Swallow-T with Swallow-C, high values of SIm were mainly observed before HG burst. Contrarily, in sensory-related PAC including Water-T with Water-C, high values of SIm were primarily observed during HG burst. The main lower frequency band indicating high values of SIm were primarily under 20 Hz. (C) Normalized SIm values are also shown. The patterns of high normalized SIm values were similar to that of high SIm values. Mouth-T, mouth triggers; Water-T, water triggers; Swallow-T, swallow triggers; Mouth-C, a mouth contacts group; Water-C, a water contacts group; Swallow-C, a swallow contacts group.

### High γ amplitude coupled with lower frequency phase

Motor-related HG activities were observed in Mouth-T with Mouth-C and Swallow-T with Swallow-C, and sensory-related HG activities were observed in Water-T with Water-C (Fig. 2A). To investigate differences between motor- and sensory-related HG activities, we used PAC analysis with the synchronization index (SI) (Cohen, 2008). We calculated the magnitude of the SI (SIm) (Fig.2B) and normalized SIm (Fig.2C) with a combination of lower frequency (5–30 Hz, 1 Hz bin) phase and fixed HG (75–150 Hz) amplitude.

In Mouth-T with Mouth-C and Swallow-T with Swallow-C, combinations where motor-related HG activities were observed, high SIm and high normalized SIm were observed before 0 s, corresponding to the time before HG power burst, rather than around 0 s, corresponding to during HG power burst. In the Water-T with Water-C, which are combinations where sensory-related HG activities were observed, high SIm values were observed after 0 s, corresponding to during HG power burst.

### Motor-related PAC appeared before HG burst, and sensory-related PAC appeared during HG burst

In motor- and sensory-related PAC, the main lower frequency band indicating high values of SIm were under 20 Hz, therefore, we fixed lower frequency range; 5-9 Hz were defined as representation of the θ band, and 10-16 Hz of the α band. With Mouth-T, Water-T, and Swallow-T, dynamic changes of SIm between θ-HG, and α-HG were plotted using Mouth-C, Water-C, and Swallow-C (Fig. 3A).

**Fig.3.**
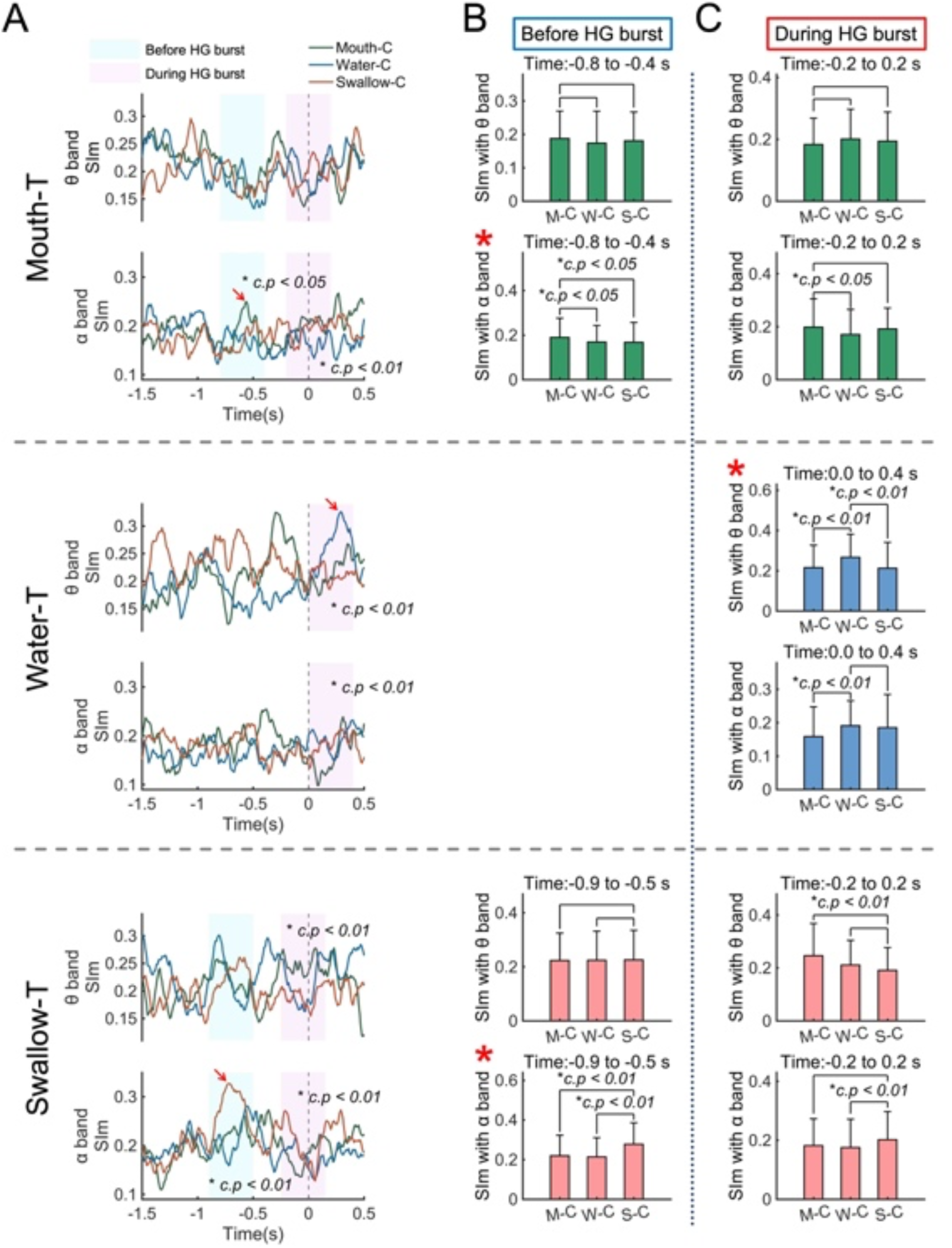
Motor-related phase-amplitude coupling (PAC) appeared before high γ (HG) burst, sensory-related PAC appeared during HG burst. (A) Dynamic changes of magnitudes of synchronization index (SIm) of θ band (5-9 Hz) and α band (10-16 Hz) coupled with HG are plotted among Mouth-T, Water-T, and Swallow-T using Mouth-C, Water-C, and Swallow-C colored by green, blue, and red respectively. The peaks of SIm are indicated by red arrows. The time of before HG burst and during HG burst are indicated as cyan mesh area and magenta mesh area in each. We compared three contacts groups using one-way analysis of variance with Bonferroni correction. (B) The SIm values calculated from each group before HG burst are compared using T-test with Bonferroni correction. In Mouth-T, α-HG SIm of Mouth-C achieved significantly higher values rather than other groups, and in Swallow-T, α-HG SIm of Swallow-C achieved significantly higher values rather than other groups (red asterisk). (C) The SIm values during HG burst were compared among three group by T-test with Bonferroni correction. In Water-T, θ-HG SIm of Water-C achieved significantly higher values rather than other groups (red asterisk). The error bars in (B) and (C) indicate standard deviation. Significant differences were indicated by black star. *c.p*, corrected *p* Mouth-T, mouth triggers; Water-T, water triggers; Swallow-T, swallow triggers; Mouth-C/M-C, a mouth contacts group; Water-C/W-C, a water contacts group; Swallow-C/S-C, a swallow contacts group.

In Mouth-T, α-HG SIm of Mouth-C showed a peak at −0.6 s which corresponded to before HG burst (Fig. 3A, top column, red arrow). Significant differences of α-HG SIm among the three contacts groups were observed before the HG burst (*c.p* < 0.05, one-way analysis of variance (ANOVA) with Bonferroni correction). In only before HG burst with α-HG SIm, Mouth-C reached significantly higher values than Water-C and Swallow-C (*c.p* < 0.05, *t*-test with Bonferroni correction; Fig. 3B and 3C, top column and red asterisk).

In Water-T, θ-HG SIm of Water-C showed a peak at 0.2 s which corresponded to during HG burst (Fig.3A, middle column, red arrow). Significant differences among the three contacts groups were observed during HG burst in θ-HG SIm (*c.p* < 0.01, ANOVA with Bonferroni correction). In θ-HG SIm during HG burst, Water-C reached the highest values compared to Mouth-C and Swallow-C (*c.p* < 0.01, T-test with Bonferroni correction; Fig. 3C, middle column and red asterisk).

In Swallow-T, α-HG SIm of Swallow-C showed a peak at −0.7 s which corresponded to before HG burst (Fig.3A, bottom column, red arrow). Significant differences among the three contacts groups were observed at before HG burst of α-HG SIm (*c.p* < 0.01, ANOVA with Bonferroni correction). In only before HG burst with α-HG SIm, Swallow-C reached significantly higher values compared to Mouth-C, and Water-C (*c.p* < 0.01, T-test with Bonferroni correction; Fig. 3B and 3C, bottom column and red asterisk).

We concluded that motor-related PAC appeared before motor-related HG burst, not during HG burst, and its main lower frequency phase was the α band, while the sensory-related PAC appeared during sensory-related HG burst, and its main lower frequency phase was the θ band (Table 2).

**Table 2.** Main low frequency band coupled with high γ amplitude.

| Event | Type | Before high $\gamma$ burst | During high $\gamma$ burst |
| --- | --- | --- | --- |
| Mouth opening | Motor | $\alpha$ band | No significant coupling |
| Swallowing | Motor | $\alpha$ band | No significant coupling |
| Water injection | Sensory | No significant coupling | $\theta$ band |

### Motor-related HG amplitude were tuned by a trough of α oscillation, and sensory-related HG amplitude were tuned by a peak of θ oscillation

We calculated the mean vectors of preferred phase of synchronization (SIp) during motor- and sensory-related SIm increasing (Fig.4A). In motor-related SIp calculated from Mouth-T with Mouth-C and Swallow-T with Swallow-C before HG burst, the angle of the mean vector ranged from 0° to 45°, however, in sensory-related SIp calculated from Water-T with Water-C during HG burst, the angle of the mean vector was approximately 270° (−90°). We observed significant nonuniformity in all groups (*c.p* < 0.01, Rayleigh test with Bonferroni correction).

**Fig.4.**
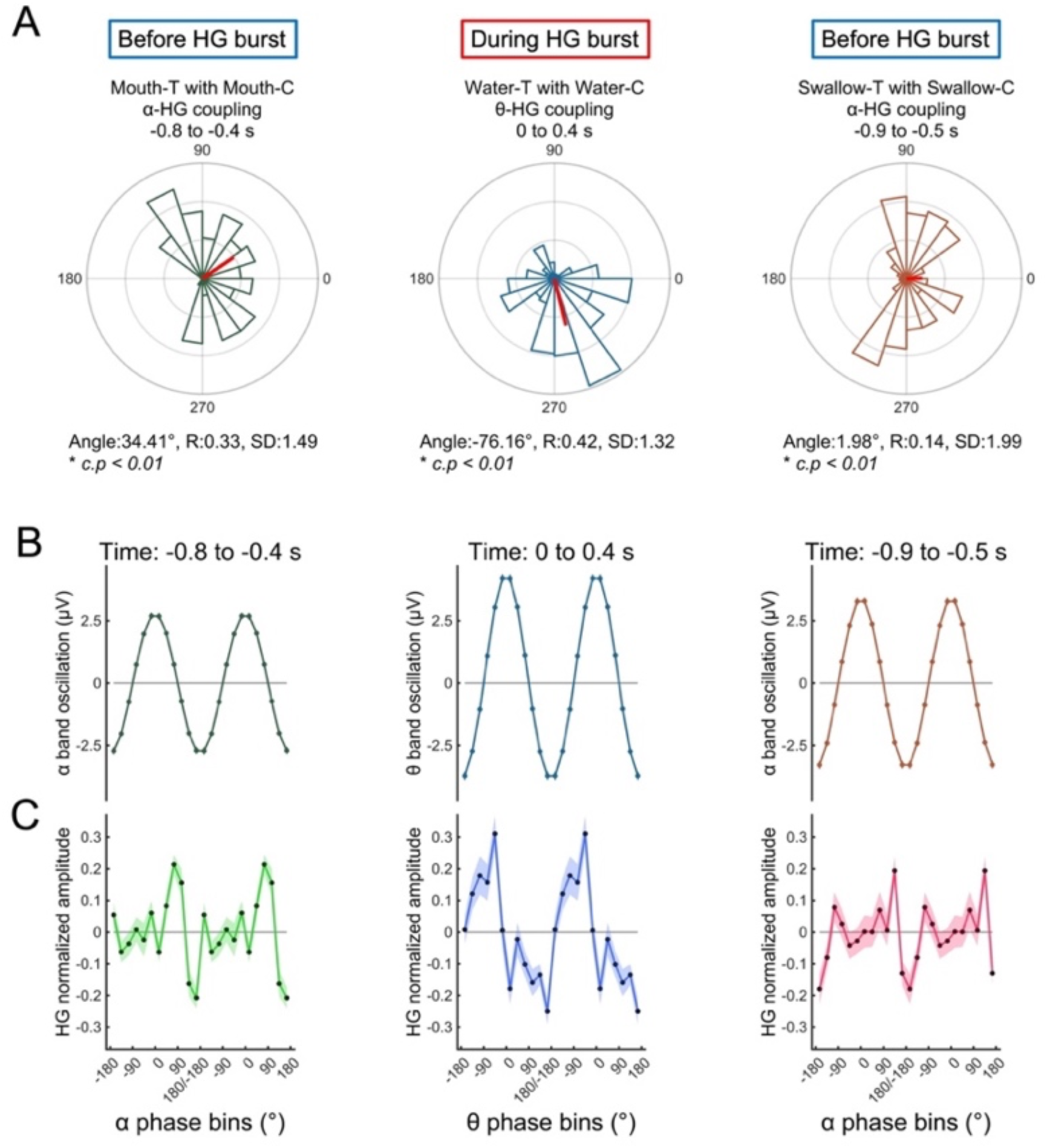
Mouth- and swallow-related high γ (HG) amplitude were tuned by the trough of α oscillations, and water-related HG amplitude were tuned by the peak of θ oscillations. (A) We calculated average values of the preferred phase of synchronization (SIp) at the time when magnitudes of synchronization index (SIm) increased. Motor-related SIm calculated from Mouth-T with Mouth-C and Swallow-T with Swallow-C increased before HG burst, and sensory-related SIm calculated from Water-T with Water-C increased during HG burst. The length, and standard deviation of the mean vector are indicated as R, and SD and corrected p-values (*c.p*) calculated by the Rayleigh test with Bonferroni correction are shown. The average angle of motor-related SIp ranged from 0° to 45°, however, in that of sensory-related SIp was approximately 270° (−90°). At the time when SIm increased, the lower frequency oscillations of lower frequency phase (B) and HG normalized amplitude tuned by lower frequency phase (C) are displayed. In mouth and swallow group, HG normalized amplitude peaked at the trough of α oscillation, and in water group, HG normalized amplitude peaked at the peak of θ oscillation. The error bars in (B) and (C) indicate 95% confidence intervals. Mouth-T, mouth triggers; Water-T, water triggers; Swallow-T, swallow triggers; Mouth-C, a mouth contacts group; Water-C, a water contacts group; Swallow-C, a swallow contacts group.

We showed the lower frequency oscillations of lower frequency phase (Fig. 4B) and HG normalized amplitude tuned by lower frequency phase (Fig. 4C) during motor- and sensory-related SIm increasing. At motor-related SIm increasing corresponding to before HG burst, HG normalized amplitude achieved the peak around the trough of α oscillations. Contrarily, at sensory-related SIm increasing corresponding to during HG burst, HG normalized amplitude peaked at the peak of θ oscillations.

There were clear differences between motor-related and sensory-related HG amplitude tuning; during SIm increase, motor-related HG amplitude were tuned by the trough of α oscillations, and sensory-related HG amplitude were tuned by the peak of θ oscillations.

### Sensory-related SIm positively correlated with HG normalized power

Finally, we investigated the correlation between SIm and HG normalized power. Using Mouth-C, Water-C, and Swallow-C, the dynamic changes of SIm and HG normalized power were displayed from before 1.5 s to after 0.5 s around each trigger (Fig. 5A and 5B). Using total implanted contacts (151 contacts), sequential correlation coefficients (r) between SIm and HG normalized power were plotted (Fig. 5C). We set the threshold of correlation coefficients that achieved 80% statistical power; the threshold was ± 0.2. At Mouth-T and Swallow-T, during SIm increased (cyan mesh area), no significant correlation was observed; however, at HG burst of Water-T (yellow mesh area), positive correlation was observed. At HG burst of Mouth-T, and Swallow-T, there was no obvious correlation.

**Fig.5.**
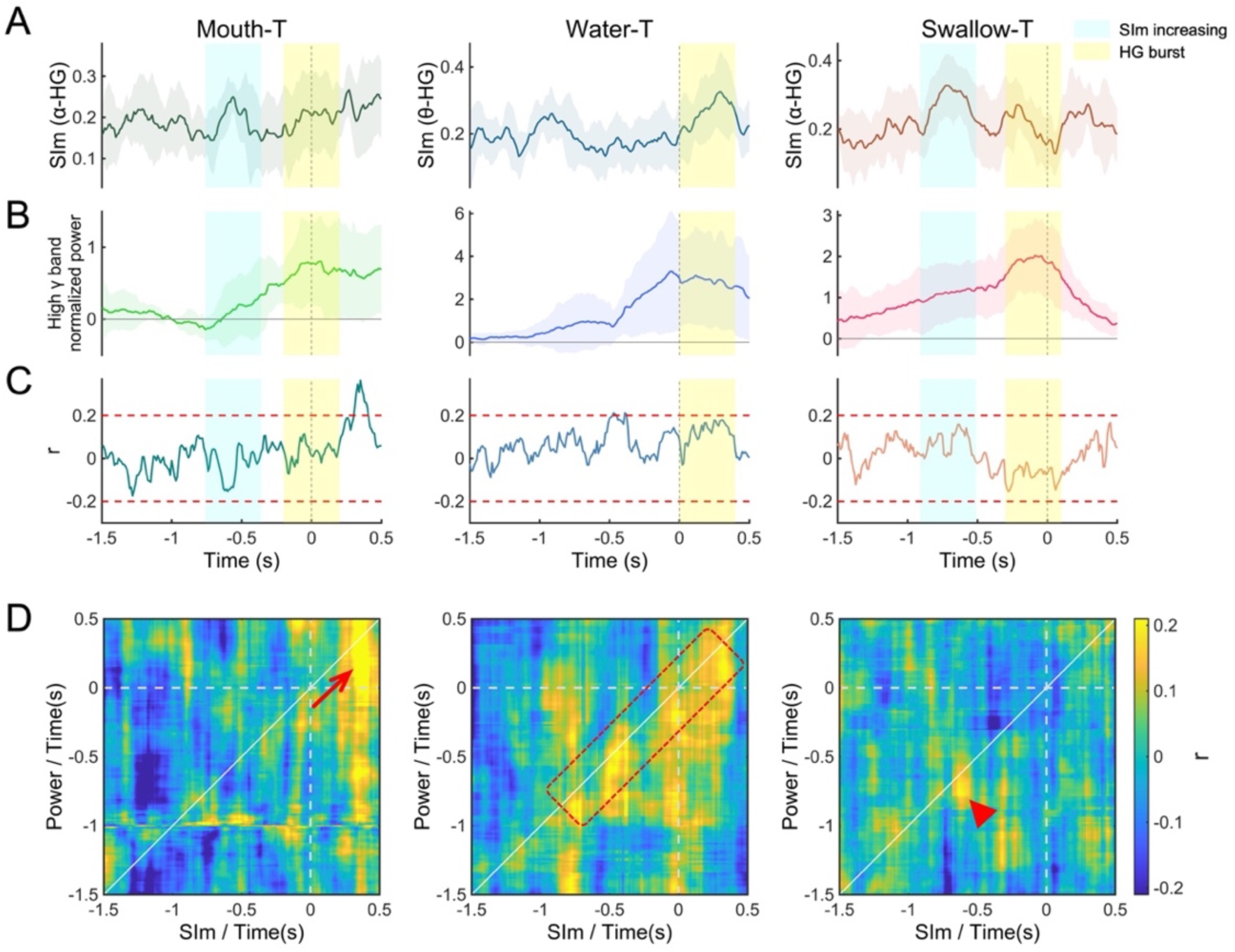
Sensory-related SIm positively correlated with HG normalized power. Using Mouth-C, Water-C, and Swallow-C, at Mouth-T, Water-T, and Swallow-T for each, dynamic changes of magnitudes of synchronization index (SIm) (A) and high γ (HG) normalized power (B) were plotted from −1.5 s to 0.5 s around each trigger. In motor-related SIm of mouth and swallow group, SIm between α band (10-16 Hz) and HG were calculated, and in sensory-related SIm of water group, SIm between θ band (5-9 Hz) and HG were also calculated. Using total 151 implanted contacts, we calculated sequential correlation coefficients (r) between SIm and HG normalized power (C). The time of SIm increasing and HG burst are indicated as cyan mesh area and yellow mesh area in each. The threshold of correlation coefficients that achieved 80% statistical power are indicated as red dashed lines (± 0.2). (D) Using total 151 implanted contacts, correlation coefficients (r) are shown for all combinations of sequential SIm (horizontal axes) and sequential HG normalized power (vertical axes), from −1.5 s to 0.5 s around each trigger. In Water-T, we observed an positive correlation along the diagonal line (red square). In Mouth-T and Swallow-T, a positive correlation was temporarily observed (red arrow and red wedge arrow. The error bars in (A) and (B) indicate 95% confidence intervals. Mouth-T, mouth triggers; Water-T, water triggers; Swallow-T, swallow triggers; Mouth-C, a mouth contacts group; Water-C, a water contacts group; Swallow-C, a swallow contacts group.

We calculated the correlation coefficients in combination with sequential SIm and HG normalized power from −1.5 s to 0.5 s around each trigger using total 151 contacts and illustrated them as a matrix (Fig. 5D). In Water-T, we observed an obvious positive correlation along the diagonal line from approximately −0.5 s to 0.5s, which corresponded to the time when HG normalized power increase (red square in Fig. 5D). The pattern that SIm and HG normalized power in the same time zone had a positive correlation was not observed in Mouth-T and Swallow-T. In Mouth-T and Swallow-T, a positive correlation was temporarily observed (red arrow and red wedge arrow in Fig.5D).

Sensory-related PAC represented by SIm showed the positive correlation with sensory-related HG activities, whereas, motor-related PAC has no obvious correlation between SIm and HG activities.

## Discussion

We used three different contacts group showing significant HG burst within each trigger (Mouth-C, Water-C, and Swallow-C). The distribution pattern of each group varied in the orofacial cortex. Mouth-C was primarily in the frontal lobe including the lateral precentral gyrus and in the ventrolateral prefrontal cortex. These regions have been reported to be actively involved in orofacial motor action (Miller et al., 2007, Loh et al., 2020). Water-C was observed in the lateral portion along the central sulcus, which were known to be activated by tactile stimulation of the buccal mucosa (Miyaji et al., 2014). Swallowing-C was observed in regions along the Sylvian fissure, that are activated by swallowing (Martin et al., 2004, Toogood et al., 2017, Satow et al., 2004, Hamdy et al., 1999, Dziewas et al., 2003, Lowell et al., 2008). The contacts’ distribution pattern of our results concordant with previous results.

A decrease in α/mu power in motor cortices is associated with in increased activation of the cortical area (Wagner et al., 2019). HG activities associated with hand movements were coupled with the α phase before the HG increase (Yanagisawa et al., 2012b), hence a hold-and-release model of HG amplitude was proposed, which indicated that strong coupling restricted the HG activities and attenuation of the coupling releases the HG activities. The high PAC state may suppress cortical information processing before movement onset, and its cessation may induce a shift to an active processing state (de Hemptinne et al., 2015a). In this study, coupling between α-HG related to mouth opening and swallowing was also observed before the motor-related HG burst. In a previous study, the maximum value of the HG amplitude associated with hand movement was locked up to the trough of the α oscillation (Yanagisawa et al., 2012b), while in our study, the HG amplitude associated with mouth opening and swallowing was locked to the trough of the α oscillation. Our finding regarding motor-related PAC was similar to that of previous studies, showing that we have successfully demonstrated the motor-specific PAC in the orofacial cortex using mouth opening and swallowing tasks.

Previous studies have demonstrated that HG activities were induced by somatosensory stimulation (Plenz and Kitai, 1996, Hashimoto et al., 1996, Hirata et al., 2002), while we showed the HG burst related to water injection. Sensory-related PAC were studied primarily using visual (Seymour et al., 2017, Voytek et al., 2010) or auditory tasks (Malinowska and Boatman-Reich, 2016, Luo and Poeppel, 2007), however, PAC studies using tasks of somatosensory stimulation are limited. In this study, during the sensory-related HG burst, the θ-HG coupling obtained high values, which indicated that the HG activities increased at the θ rhythm. The HG amplitude coupled with θ phase related to water injection was locked to the peak of the θ oscillation, which are the contrary to motor-related PAC. We conclude that the differences in the time when coupling occurs, in the lower frequency band which coupled with HG amplitude, and in the lower frequency phase which modulates HG amplitude between motor output and somatosensory input may reflect the differences between motor- and sensory-related neural processing.

The preferred low-frequency coupling rhythm modulating HG amplitude could be altered by task demands (Voytek et al., 2010). δ-HG coupling is associated with auditory and visual selective-attention task (Lakatos et al., 2008), while θ-HG coupling is associated with a auditory task (Malinowska and Boatman-Reich, 2016, Luo and Poeppel, 2007, Canolty et al., 2006), a working memory (Axmacher et al., 2010), and an episodic memory (Heusser et al., 2016); an α-HG coupling is associated with a visual task (Voytek et al., 2010, Seymour et al., 2017), and motor task (Yanagisawa et al., 2012b). The regions where PAC were observed also varied among the tasks; the auditory cortex with auditory task (Malinowska and Boatman-Reich, 2016), the visual/occipital cortex with a visual task (Seymour et al., 2017, Voytek et al., 2010), and the sensorimotor cortex with a motor task (Yanagisawa et al., 2012b). In addition to these findings, our results also showed that each of θ-HG and α-HG coupling were predominant in the orofacial cortex during a sensory- and a motor-task respectively.

HG power is strongly correlated with neural firing rate (Ray et al., 2008), and neural spiking is locked to the trough of the α oscillations (Haegens et al., 2011). Therefore, during mouth opening- and swallowing-related α-HG coupling occurred, the HG amplitude might be modulated by the trough of the α oscillations. Moreover, it has been proposed that α oscillations represented a pulsed-inhibition of ongoing neural activity (Mathewson et al., 2011), and therefore, it may be reasonable that our results showed no HG activities during appearance of motor-related α-HG coupling.

EEG studies suggest that θ oscillations are related to both sensory and motor behaviors (Tomassini et al., 2017), and a general property of brain activity (Ramayya et al., 2020). A θ-HG coupling was observed not only in the cortex but also the hippocampus (Tort et al., 2008). Animal experiments showed that in the epileptic brain, the hippocampal θ rhythm was increased (Kitchigina and Butuzova, 2009), and seizure-related HFA were coupled with θ band (Hashimoto et al., 2020a, Hashimoto et al., 2020b). The θ oscillations are involved in both of sensory and motor neural processing, and both of physiological and pathological neural processing, and in this study, we provide a new insight into the sensory-related θ-HG coupling.

Epilepsy studies reported a positive correlation between coupling values and high frequency amplitude (Weiss et al., 2016, Hashimoto et al., 2020b). We showed the continuous positive correlation between the θ-HG coupling and HG power related to sensory stimulation and inferred that a tactile stimulation by water-injection into the oral cavity induced the HG activities in the lateral central sulcus, while the HG activities were increased at the θ rhythm. However, the causality that coupling induced HG power or HG power induced coupling remains unclear. We expected that motor-related α-HG coupling might correlate positively with later HG power during motor-related HG burst occurred. However, contrary to our expectation, no obvious positive correlation between coupling and HG power was found; nonetheless, a positive correlation was temporally observed at approximately 0.3 s of Mouth-T (Fig. 5D, red arrow). We inferred that this positive correlation was caused by sensory input since water-bolus were injected immediately after mouth-opening.

In conclusion, using Mouth-T and Swallow-T as a representative of motor-related task and Water-T as a representative of sensory-related task in combination with three different contacts groups plotted over the orofacial cortex, we demonstrated the differences in α-HG coupling appeared before motor-related HG power burst and θ-HG coupling appeared during sensory-related HG burst. Moreover, we showed that during appearance of low and high frequency coupling, motor-related HG amplitudes were modulated at the trough of α oscillations, and sensory-related HG amplitudes were modulated at the peak of θ oscillations. These neuro-oscillatory differences may reflect the differences between motor- and sensory-related neural processing in the cortex.

## Limitations of the Study

First, we focused only the orofacial cortex; multiple cortices are activated during swallowing (Ertekin and Aydogdu, 2003), however our study could not demonstrate cortical activities other than in orofacial region. Second, in this study, we divided swallowing events into three parts (mouth opening, water injection, and swallowing). The time of mouth opening and water injection were detected using the video captured by Kinect v2, having 30 frames per second, therefore, a maximum gap of 33 msec may occur. For more precise analysis, we must improve the detection methods of the time of mouth opening and water injection. Third, we defined θ band as 5-9 Hz and α band as 10-16 Hz. However, the frequency range generally used are 4-8 Hz, for θ band, (Canolty et al., 2006) and 8-14 Hz, for α band (Haegens et al., 2011). We set the frequency range in an effort to demonstrate that the frequency range in which coupling occurred related to motor- and sensory-related task differ. Finally, we focused only on frequencies > 5 Hz. Since we used a 400 ms time-window for SIm calculation, the minimum frequency in which at least two oscillation cycles were included was 5 Hz (Cohen, 2008).

## Resource Availability

### Lead Contact

Further information and requests for resources should be directed to and will be fulfilled by the Lead Contact, Masayuki Hirata.

### Materials Availability

This study did not generate new unique reagents or materials.

### Data and Code Availability

Original/source data in the paper is available upon request.

## Transparent Methods

### Participants

Eight patients with intractable temporal lobe epilepsy participated in this study (four females, 10’s–50’s years of age) (Table 1). They were admitted to Osaka University Hospital from April 2015 to July 2019, and underwent intracranial electrodes placement for intracranial electroencephalogram (iEEG) study. They had normal swallowing function, which was confirmed by medical interview. All participants or their guardians were informed of the purpose and possible consequences of this study, and written informed consent was obtained. The Ethics Committee of Osaka University Hospital approved this study (Nos. 08061, 16469).

### Intracranial electrodes

Different types of electrodes (Unique Medical Co. Ltd., Tokyo, Japan), including grid, strip, and depth types, were implanted in the subdural space during conventional craniotomy as a clinical epileptic surgery. For analysis, we chose planar-surface platinum grid electrodes (4×5 contacts array) that were placed over the lateral portion of the central sulcus corresponding to the orofacial cortex. The number of total implanted contacts and selected contacts in each participant are shown in Table 1. The diameter of the contacts was 3 or 5 mm, and the intercontact center-to-center distance was 5, 7, or 10 mm.

### Electrode location

Preoperative structural MRI was obtained with a 1.5-T or 3.0-T MRI scanner, and postoperative computed tomography (CT) scans were acquired with the implanted electrodes in place. The implanted electrodes that were obtained from the CT images were overlaid onto the 3-dimensional brain renderings from the MRI volume that was created by FreeSurfer software (https://surfer.nmr.mgh.harvard.edu). We obtained the Montreal Neurological Institute (MNI) coordinates of the implanted contacts with Brainstorm software (http://neuroimage.usc.edu/brainstorm/), which merged the preoperative MRI scans and postoperative CT scans. The location of the implanted electrodes that was visualized by Brainstorm was then confirmed by intraoperative photographs.

### Task

The experiments were performed approximately 1 week after surgical electrode placement when all participants fully recovered from surgery. The participants were asked to sit on a chair and to remain still, especially without moving their mouth. The participants then opened their mouths, and the examiner injected 2 mL of water into their mouths with a syringe. We requested that the participants swallow it at their own pace without external cueing to prevent erroneous volitional water swallowing (aspiration). After we confirmed that the participants had completed one swallowing movement, the next water bolus was injected.

### Swallowing monitoring

For noninvasive swallowing detection, we used an electroglottograph (EGG), a microphone, and a motion-tracking system (Supplementary Fig.1A). A laryngograph (Laryngograph Ltd, London, UK) was used as an EGG and recorded the neck impedance changes associated with swallowing (Firmin et al., 1997) (Supplementary Fig.1B). A pair of contacts was placed on the neck skin below the thyroid cartilage of the participants at an intercontact center-to-center distance of 25 mm and was held in place by an elastic band.

Sounds of swallowing due to the bolus passing through the pharynx were detected by a throat microphone (Kusuhara et al., 2004) (Supplementary Fig.1C). We connected the throat microphone (Inkou mike; SH-12iK, NANZU, Shizuoka, Japan) to the laryngograph to record the swallowing sounds. The shape of the microphone was arched to fit around the participant’s neck. The sampling rate of a laryngograph and a throat microphone was 24,000 Hz.

We captured the motion of the participants at 30 frames per second with the motion-tracking system, which was newly developed by us using Kinect v2 (Microsoft, Redmond, Washington, USA) (Hashimoto et al., 2018). The participants were seated facing Kinect v2, which was placed on a tripod at a distance of one meter, and their mouth and throat movements were captured automatically.

An electric stimulator (NS-101; Unique Medical, Tokyo, Japan) supplied digital synchronizing signals to the laryngograph and a 128-channel digital EEG system. The signals made an LED light flash, which was captured by the RGB camera of Kinect v2. The digital triggers and LED light flash enabled us to synchronize the multimodal data of an EGG, a microphone, the video captured by the RGB camera, and an intracranial EEG. The multimodal data enabled us to noninvasively monitor the swallowing movements. The video captured by the RGB camera enabled us to the detect time when the participants opened their mouths and when the water bolus was injected into the mouth.

### Signal segmentation based on swallowing-related events

The swallowing onset time was determined at the time when the impedance waveform reached the peak (Supplementary Fig.1B). Swallowing sounds occurred frequently as the bolus of water passed through the pharynx (Kusuhara et al., 2004), and their evaluation in conjunction with the EGG helped us to judge whether the impedance change was caused by swallowing (Supplementary Fig.1C). Additionally, we confirmed that the changes in impedance and sounds corresponded to water swallowing by using the video captured by the RGB camera. We inserted swallowing triggers, which corresponded to the swallowing onset time, into the iEEG data.

We could also detect the time when the participant opened his/her mouth and when the examiner injected water into the participants’ mouth using the video captured by the RGB camera. Mouth triggers and water triggers, which corresponded to different times, were also inserted into iEEG data (the number of each trigger is shown in Table 1).

### Data acquisition and preprocessing

The iEEG signals were measured with a 128-channel digital EEG system (EEG 2000; Nihon Kohden Corporation, Tokyo, Japan) and digitized at a sampling rate of 1000 Hz with a 0.3 to 333 Hz bandpass filter to prevent aliasing, and a 60-Hz notch filter to eliminate the AC line noise. Before any further processing, contacts containing external noise or epileptic discharge were excluded from further analyses. The iEEG signals were digitally re-referenced to a common average of implanted total contacts in each participant. We analyzed the iEEG signals that were time-locked to the triggers.

Throughout the following analyses, a bandpass filter using a two-way least-squares finite impulse response filter (pop_eegfiltnew.m from the EEGLAB version 14.1.2b, https://sccn.ucsd.edu/eeglab/index.php) was applied to the iEEG signals. To prevent edge-effect artifacts, additional 250 msec data remained at the initial and end points of the pre band-pass filtered signals. After band-passed filtering, the 250 msec extra data were excluded.

### High γ power contour map

To create power contour maps, the power of each contact was constructed from the preprocessed iEEG signal by using a band-pass filter of high γ band (75–150 Hz) in combination with the Hilbert transformation (Cohen, 2008). We calculated the averaged power during 0.5 s time-window, and the averaged power was normalized with the mean and standard deviation of the power during −5.0 to −4.5 s of the swallowing triggers.

### Extraction of contacts

Significant high γ power increasing contacts (single-sided permutation tests with Bonferroni correction) were plotted over the left hemisphere of MNI normalized brain using Brainstorm software. The contacts attached to the right hemisphere were transposed to the left hemisphere.

The contacts indicating the maximum values of the significant high γ power increase in the mouth, water, and swallow triggers were extracted from each participant; in total, eight contacts within each trigger were plotted on the left hemisphere of MNI normalized brain. These group of contacts associated with each trigger were defined as mouth-related contacts (Mouth-C), water-related contacts (Water-C), and swallowing-related contacts (Swallow-C).

### Dynamic frequency power changes

We obtained averaged power waveforms of Mouth-C, Water-C, and Swallow-C relative to each trigger (mouth triggers, Mouth T; water triggers, Water T; swallowing triggers, Swallow T) in high γ band. A power time series were constructed from a band-pass filtered signal in combination with the Hilbert transformation (Cohen, 2008). The power time series were normalized by the power from base-time (−1.0 s to −0.9 s of Mouth T). Averaged frequency power values were calculated from a 100 or 400 msec time-window which was shifted every 10 msec.

### Phase amplitude coupling analyses

A synchronization index (SI) (Cohen, 2008) was used to measure the strength of coupling between the high γ amplitude and lower frequency phase. Hilbert transformation was performed on the bandpass-filtered signals to obtain complex-valued analytic signals [Z(t)]. The amplitude [A(t)] and phase [φ(t)] were calculated from the complex-valued signals using Equation 1.

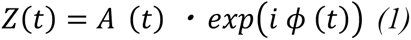

The lower frequency phase was calculated using the angle of the Hilbert transformation in the bandpass-filtered signal. The center of a bandpass filter was shifted every 1 Hz from 5 to 30 Hz, and the width of bandpass filter were ± 2.5 Hz. The high γ amplitude was calculated using the squared magnitude of the Hilbert transformation in the 75–150 Hz bandpass filtered signal. Then, the phase of this amplitude was computed using the Hilbert transformation. The SI was calculated using Equation 2.

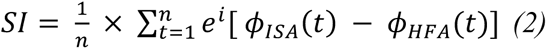

where “n” is the number of data points. We used a 400 ms time-window at a sampling rate of 1,000 Hz, therefore, the “n” value was 400. Since we used a 400 ms time-window, the minimum frequency in which at least two oscillation cycles were included was 5 Hz.

Since SI is a complex number, we used the magnitude of SI (SIm) in our calculations. SIm varies between 0 and 1, with 0 indicating completely desynchronized phases and 1 indicating perfectly synchronized phases.

The preferred phase of synchronization (SIp) was calculated by the arctan (image [SI] / real [SI]). SIp varies between −180° and +180°.

### Normalized SIm

The boot-strapping technique was used (Cohen, 2008). The phase time series of high frequency power amplitude is shifted in time by some random amount, and boot-strapping SIm (SImb) is calculated with the lower frequency phase. This procedure was repeated 1,000 times to create a distribution of SImb. The normalized SIm was calculated using the mean and standard deviation of the distribution of SImb.

### Phase-conditioned analysis

We calculated the mean vector (angle and length) using the CircStat toolbox(Berens, 2009). To identify the lower frequency phase to which the high γ amplitude was coupled, we calculated the average oscillations of lower frequency band and the normalized amplitude of high γ band within each lower frequency band phase bin of 30°: −180° – −150°, −150° – −120°, …, and 150° – 180°.

### Correlation analysis

Using all the implanted contacts (total 151 contacts), we calculated Pearson correlation coefficients between SIm and the high γ normalized power. The values greater than +3 standard deviation (SD) or less than −3 SD were excluded as outliers. Using Monte Carlo method, we set the threshold of correlation coefficients that achieved 80% statistical power.

### Statistics

For statistical evaluation of high γ power changes, we used a permutation test (Maris and Oostenveld, 2007). For comparison of two groups, T–test was used. For correction of multiple comparisons, we used Bonferroni correction. For comparison of three groups of mouth triggers, water triggers, and swallowing triggers, we used one-way analysis of variance (ANOVA). We performed the Rayleigh test to evaluate the nonuniformity of SIp using the CircStat toolbox(Berens, 2009).

## Acknowledgments

This work was supported by Japan Society for the Promotion of Science (JSPS) KAKENHI [Grant nos. JP26282165(Masayuki Hirata), JP18H04166(Masayuki Hirata), JP18K18366(Hiroaki Hashimoto)], by the Ministry of Internal Affairs and Communications (Masayuki Hirata), by a grant from the National Institute of Information and Communications Technology (NICT) (Masayuki Hirata), and by a grant from the National Institute of Dental and Craniofacial Research (NIDCR)-RO1 DE023816 (Kazutaka Takahashi).

## Author contributions

M.H. designed the study. H.H. performed the experiments, assisted with the epileptic medical treatment, created the MATLAB program and analyzed the data, created all the figures and tables, and was primarily responsible for writing the manuscript. F.Y. H.M. and M.H. assisted in the acquisition of the measurements. S.K. developed some devices to help with the measurements. H.K., S.O., N.T., H.M.K. and M.H. performed the epilepsy surgery, and Ta.Ya. assisted with the epileptic medical treatment. M.H., K.T. and Ta.Ya advised H.H. on scientific matters. M.H. and K.T. revised the manuscript. To.Yo., H.K. and M.H. supervised the experiments and analyses. All authors reviewed the manuscript.

## Supplementary Information

**Supplementary Fig 1.**
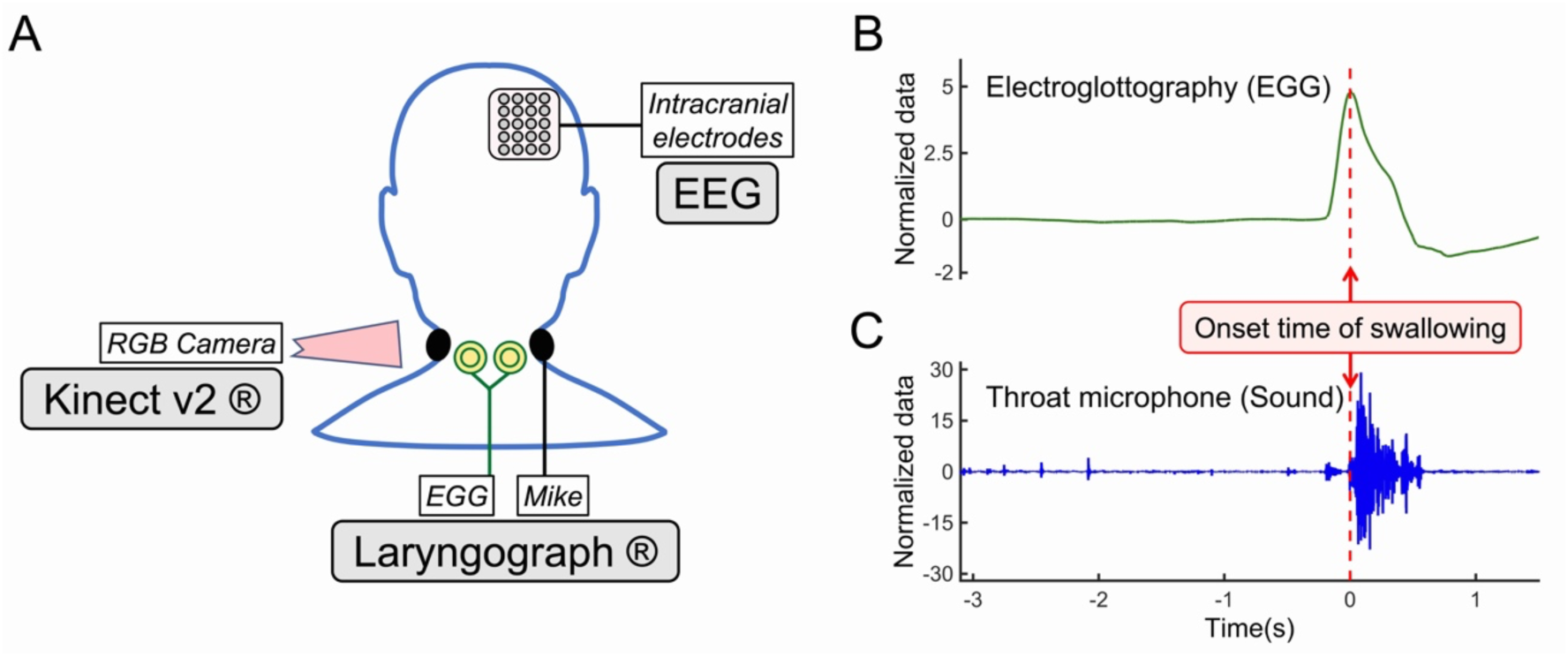
Swallowing monitoring. Intracranial EEG were recorded during participants swallowed. The swallowing was monitored by an RGB camera of Kinect v2 (Microsoft), and an electroglottograph (EGG) and a microphone of Laryngograph (A). Across-trials averaged impedance waveforms of an EGG (B) and a throat microphone (sound) (C), from Participant-1 (P1) are shown. For analysis, the onset of swallowing was defined as the peak time of an impedance waveform.

